# Thiostrepton hijacks pyoverdine receptors to inhibit growth of *Pseudomonas aeruginosa*

**DOI:** 10.1101/567990

**Authors:** Michael R. Ranieri, Derek C. K. Chan, Luke Yaeger, Madeleine Rudolph, Sawyer Karabelas-Pittman, Hamdi Abdo, Jessica Chee, Hanjeong Harvey, Uyen Nguyen, Lori L. Burrows

**Affiliations:** Department of Biochemistry and Biomedical Sciences, and the Michael G. DeGroote Institute for Infectious Diseases Research, McMaster University, Hamilton, ON

**Author notes:** **Correspondence to:** Dr. Lori L. Burrows, 4H18 Health Sciences Centre, 1280 Main St. West, Hamilton, ON L8S 4K1 Canada.

**Keywords:** antibiotic, thiopeptide, iron, siderophore, gram negative

## Abstract

*Pseudomonas aeruginosa* is a biofilm-forming opportunistic pathogen and intrinsically resistant to many antibiotics. In a high-throughput screen for molecules that modulate biofilm formation, we discovered that the thiopeptide antibiotic, thiostrepton (TS) - considered inactive against Gram-negative bacteria - stimulated *P. aeruginosa* biofilm formation in a dose-dependent manner. This phenotype is characteristic of exposure to antimicrobial compounds at sub-inhibitory concentrations, suggesting that TS was active against *P. aeruginosa*. Supporting this observation, TS inhibited growth of a panel of 96 multidrug-resistant (MDR) *P. aeruginosa* clinical isolates at low micromolar concentrations. TS also had activity against *Acinetobacter baumannii* clinical isolates. Expression of Tsr - a 23S rRNA-modifying methyltransferase - in trans conferred TS resistance, confirming that the drug acted via its canonical mode of action, inhibition of ribosome function. Deletion of oligopeptide permease systems used by other peptide antibiotics for uptake failed confer TS resistance. TS susceptibility was inversely proportional to iron availability, suggesting that TS exploits uptake pathways whose expression is increased under iron starvation. Consistent with this finding, TS activity against *P. aeruginosa* and *A. baumannii* was potentiated by FDA-approved iron chelators deferiprone and deferasirox. Screening of *P. aeruginosa* mutants for TS resistance revealed that it exploits pyoverdine receptors FpvA and FpvB to cross the outer membrane. Our data show that the biofilm stimulation phenotype can reveal cryptic sub-inhibitory antibiotic activity, and that TS has activity against select multidrug resistant Gram-negative pathogens under iron-limited growth conditions, similar to those encountered at sites of infection.

## INTRODUCTION

Bacterial pathogens are rapidly evolving resistance to available antibiotics, creating an urgent need for new therapies. Gram-negative bacteria are particularly challenging to treat because their outer membranes limit the access of many drugs to intracellular targets (1). Resistance arises when bacteria accumulate target mutations, acquire specific resistance determinants, increase drug efflux, and/or enter antibiotic-tolerant dormant or biofilm modes of growth (2). Biofilms consist of surface-associated bacteria surrounded by self-produced extracellular polymeric substances (EPS). Biofilm architecture allows for development of phenotypic heterogeneity that leads to variations in susceptibility as well as the formation of drug-tolerant persister cells (3). Approaches with the potential to preserve current antibiotics include combining them with biofilm inhibitors, resistance blockers (e.g. ampicillin with clavulanic acid or piperacillin with tazobactam), efflux inhibitors (e.g. PAβN), outer membrane permeabilizers, or coupling them to molecules such as siderophores that are actively imported, so-called Trojan horses (4).

Among the bacterial pathogens deemed most problematic by the World Health Organization is the Gram-negative opportunist, *Pseudomonas aeruginosa* (5). It infects immunocompromised patients – particularly those with medical devices – and is a major problem for people with severe burns or cystic fibrosis (6). It is intrinsically resistant to many antibiotics and readily forms biofilms, further enhancing its ability to evade therapy (7). The low permeability of its outer membrane and expression of multiple efflux pumps that extrude a wide variety of substrates, coupled with its propensity to form biofilms, limits the repertoire of effective *anti-Pseudomonas* antibiotics (8–10). Here with the initial aim of identifying potential modulators of *P. aeruginosa* biofilm formation, we screened a collection of bioactive molecules including previously FDA-approved off-patent drugs. During this work, we identified several molecules that stimulated biofilm formation beyond our arbitrary cutoff of 200% of the vehicle control. Investigation of one such stimulatory compound, thiostrepton (TS), revealed that it had low micromolar activity against *P. aeruginosa* in minimal medium. Through a series of investigations, we showed that TS gains access to its ribosomal targets by exploiting iron limitation-dependent uptake pathways. These data show that the biofilm stimulation phenotype can reveal cryptic antibiotic activity when concentrations are too low (or growth conditions not conducive) to inhibit growth.

## RESULTS

### Thiostrepton stimulates *P. aeruginosa* biofilm formation

We used a previously described *P. aeruginosa* biofilm assay (11) to screen a bespoke collection of 3921 bioactive molecules that includes ~1100 FDA-approved, off-patent drugs and antibiotics (12). The molecules were screened in duplicate at 10 μM in a dilute growth medium consisting of 10% lysogeny broth (LB), 90% phosphate buffered saline (henceforth, 10:90) to identify molecules capable of modulating biofilm formation. This medium was chosen to minimize the amount of biofilm formed in the presence of the vehicle control, so that molecules that stimulated biofilm formation could be more easily identified. The hits were divided into planktonic growth inhibitors (60 compounds), biofilm inhibitors (defined as those resulting in ≤50% of vehicle control biofilm, 8 compounds), or biofilm stimulators (those resulting in ≥200% of vehicle-treated control biofilm, 60 compounds) (**Supplementary Table S1**). The hit rate of ~3% was relatively high for a primary screen, but all the molecules in this curated collection have biological activity. The hits belonged to a variety of chemical classes and included drugs with nominally eukaryotic targets.

Multiple studies showed that sub-inhibitory concentrations of antibiotics from a variety of classes and with different mechanisms of action (MOA) stimulate *P. aeruginosa* biofilm formation, although the specific pathways underlying this response remain unclear (13–18). Among the molecules in our screen that stimulated biofilm formation was the thiopeptide antibiotic, thiostrepton (TS; **Fig 1A**). This response intrigued us because TS is considered ineffective against Gram-negative bacteria due to the impermeability of the outer membrane (OM) to large hydrophobic compounds (19, 20). In dose-response experiments in 10:90 medium, biofilm levels increased while planktonic cell density decreased with increasing TS concentrations to 10 μM (17 μg/ml), the maximum that could be tested due to its limited solubility (**Fig 1B**).

**Figure 1.**
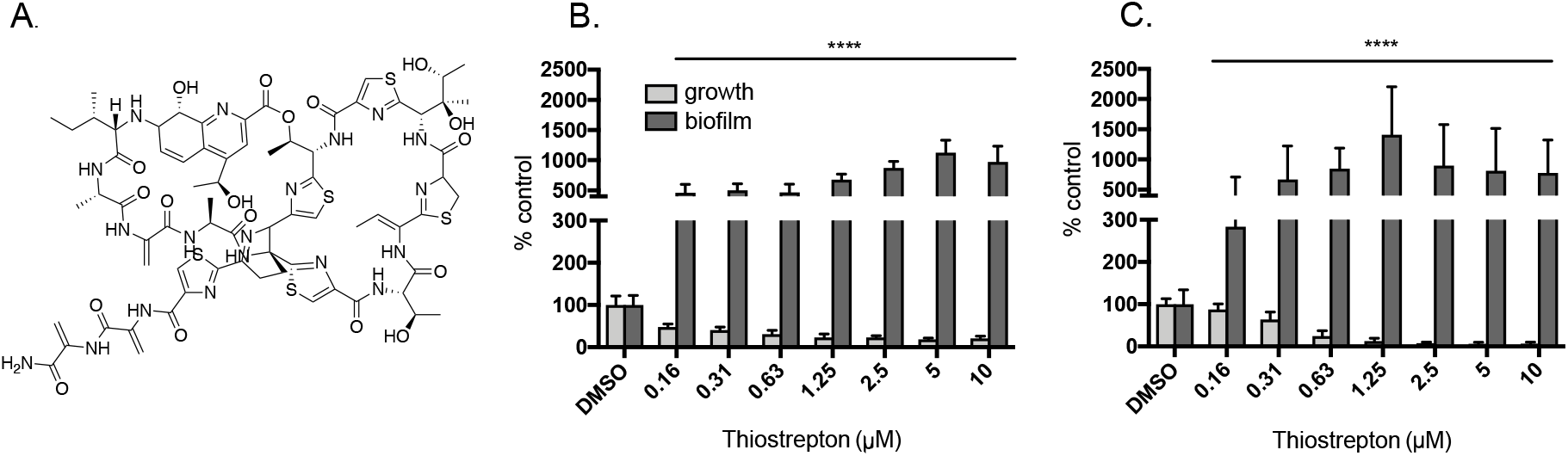
Thiostrepton stimulates *P. aeruginosa* biofilm formation. **A.** Structure of thiostrepton (TS). **B.** TS stimulated biofilm formation (absorbance of eluted crystal violet at 600 nm, plotted as percent of the DMSO control) of *P. aeruginosa* PAO1 and decreased planktonic cell density (optical density at 600 nm, plotted as percent of the DMSO control) in 10:90 medium in a dose-dependent manner, up to its maximum soluble concentration of 10 μM (17 μg/ml). **C.** In VBMM, PAO1 biofilm formation was stimulated by TS, while planktonic cell density decreased below the level of detection at concentrations above 1.25 μM. Y-axes are split to better display growth values. Assays were performed at least 3 times in triplicate. **** p<0.0001

### Growth in minimal media increases susceptibility of *P. aeruginosa* to TS

Environmental conditions can modulate the expression or essentiality of antibiotic targets or alter the availability of particular nutrients (21), leading to changes in susceptibility. We hypothesized that the biofilm response of *P. aeruginosa* to TS may be the result of nutrient deficiency in 10:90, which was more limiting to *P. aeruginosa* growth than M9 minimal medium (**Supplementary Fig S1**). Growth in Vogel Bonner Minimal Media (VBMM) in the absence of TS was similar to that in 10:90 (**Supplementary Fig S1**) but in the presence of TS, planktonic cell density decreased to below the level of detection at concentrations above ~1.25μM (**Fig 1C**). These data suggested that nutrient limitation enhances susceptibility of *P. aeruginosa* to TS.

### The ribosomal methyltransferase Tsr protects *P. aeruginosa* against TS

The established MOA for TS is inhibition of protein translation through direct binding to bacterial ribosomes (22). However, because TS also has anti-parasitic and anti-neoplastic activities (23, 24) we considered the possibility that it might inhibit *P. aeruginosa* growth in a novel way. To validate the MOA, we expressed a resistance gene, *tsr*, from a plasmid in *P. aeruginosa* strains PAO1 and PA14. *tsr* encodes a 23s rRNA methyltransferase, used by TS producer *Streptomyces azureus* to prevent self-intoxication (25). Tsr methylates the conserved A1067 residue of 23s rRNA, impairing binding of TS to its target (26). Expression of *tsr* in trans increased TS resistance of both PAO1 and PA14 compared to vector-only controls (**Fig 2**). PAO1 was resistant up to the maximum soluble TS concentration of 10 μM, while resistance of PA14 was significantly increased compared to control, although not to the same extent as PAO1. These results suggest that TS inhibits growth via its canonical MOA of ribosome binding, implying that it can cross the *P. aeruginosa* OM to access the bacterial cytoplasm.

**Figure 2.**
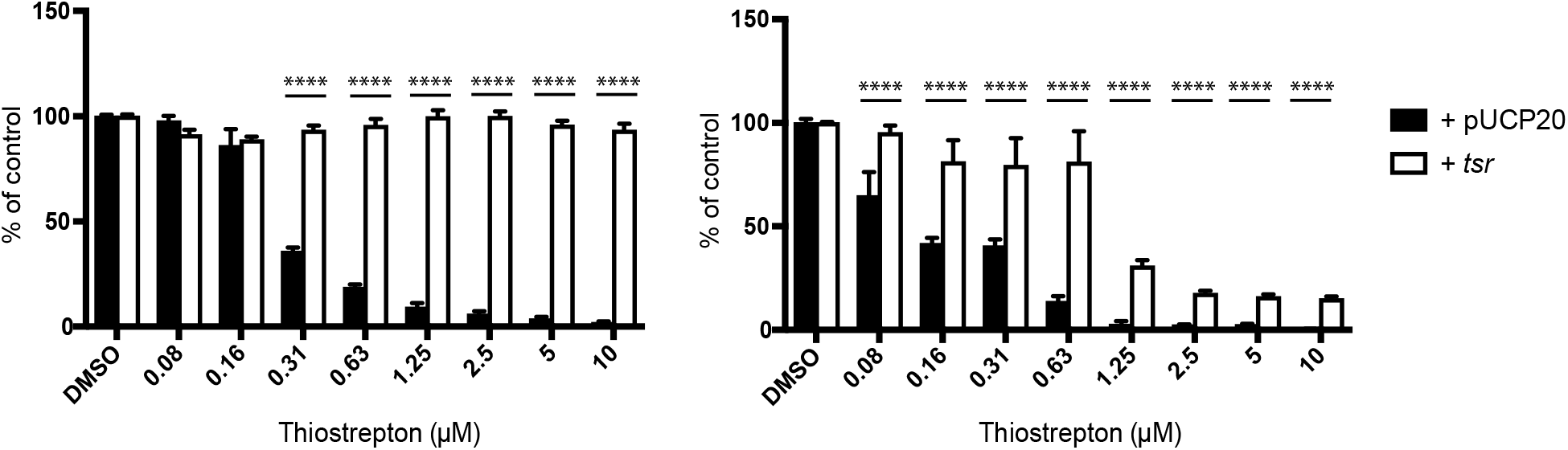
Expression of Tsr in trans reduces susceptibility of *P. aeruginosa* to thiostrepton. Expression of the *tsr* gene from *Streptomyces azureus* in trans from pUCP20 in two strains of *P. aeruginosa* reduced their susceptibility to TS in VBMM, suggesting it inhibits growth via its canonical mode of action, disrupting translation. **A.** Growth of PAO1 (OD_600_ plotted as percent of the DMSO control); **B.** Growth of PA14. Each assay was performed at least 3 times in triplicate. **** p<0.0001

### TS susceptibility increases in the presence of iron chelators

To understand the reason for increased TS susceptibility of *P. aeruginosa* in VBMM compared to 10:90, we considered the differences in nutrient availability between the two media types. The primary carbon source in 10% LB is amino acids (27) while the carbon source in VBMM is citrate (28). Citrate can chelate divalent cations including calcium and magnesium, which are important for OM integrity. We hypothesized that this chelation effect may increase OM permeability. To stabilize the OM, we repeated the dose response assay in VBMM supplemented with 100 mM MgCl2 but saw no effect on susceptibility (**Supplementary Fig S2A**). Since TS is a thiopeptide, we next hypothesized that amino acid limitation during growth in VBMM may increase uptake of TS, leading to growth inhibition. To test this, we supplemented VBMM with 0.1% casamino acids, but saw no change in TS susceptibility (**Supplementary Fig 2B**). Further, simultaneous deletion of components of the Opp (Npp) peptide transport system, exploited by other peptide antibiotics for entry (29, 30), and a homologous system, Spp, had no effect on TS susceptibility (**Supplementary Fig S2C**).

We next considered that VBMM was more iron-limited than 10:90, which contains trace iron from yeast extract and peptone. Under iron limitation, bacteria secrete siderophores into the extracellular milieu to scavenge the metal. Specialized receptors then transport siderophore-iron complexes back into the cell. Some antimicrobials, including sideromycins, pyocins, and bacteriocins, use siderophore receptors to access intracellular targets (31–34), and we hypothesized that TS may use this strategy. We compared *P. aeruginosa* PAO1 grown in 10:90 with increasing concentrations of TS alone (**Fig 3A**) or with 0.1μM EDDHA, a membrane-impermeable iron chelator (35) (**Fig 3B**). Addition of EDDHA shifted biofilm stimulation and growth inhibition to lower concentrations of TS compared to 10:90 alone. Supplementation of 10:90 plus 0.1μM EDDHA with 100μM FeCl_3_ increased planktonic cell density and reduced biofilm stimulation (**Fig 3C**). These data suggest that TS susceptibility is inversely proportional to iron availability, and that TS may exploit siderophore receptors to cross the OM of *P. aeruginosa*.

**Figure 3.**
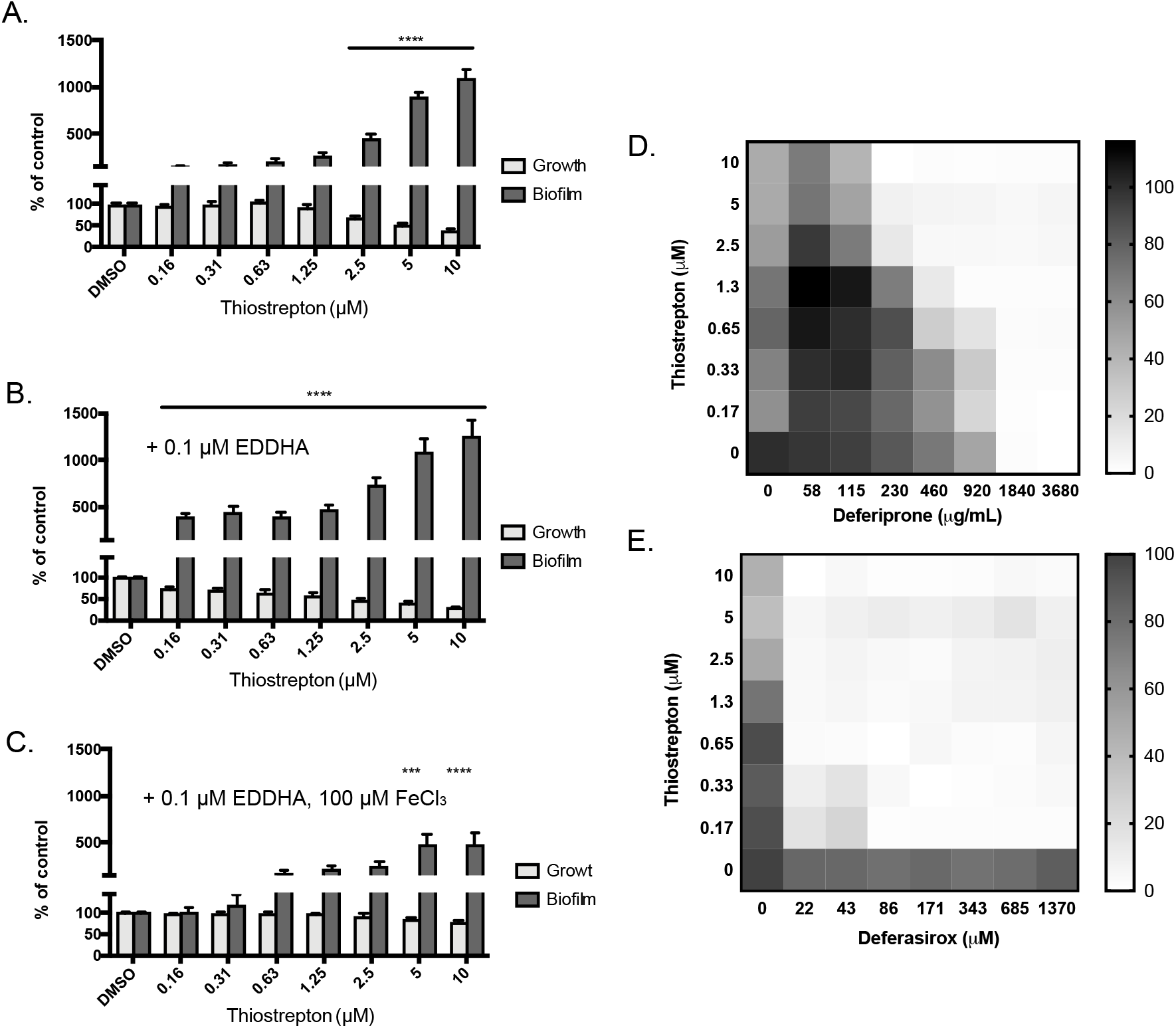
Thiostrepton activity is potentiated by iron chelators. Biofilm stimulation by TS in 10:90 medium increased in the presence of 0.1 μM EDDHA, a cell-impermeant iron chelator, while further addition of 100 μM FeCl_3_ increased the concentration of TS required for biofilm stimulation and growth inhibition. **A.** PAO1 growth (OD_600_) and biofilm (absorbance of CV at 600 nm) in 10:90 medium alone; **B.** 10:90 plus 0.1μM EDDHA; **C.** 10:90 plus 0.1μM EDDHA and 100 μM FeCl_3_. Checkerboard assays plotted as percent growth of the DMSO control (0,0 μM at lower left) showed that TS activity against PAO1 was potentiated by FDA-approved iron chelators, **D.** deferiprone and **E.** deferasirox. The highest concentrations of DFP (3680 μM) and DSX (1370 μM) are each equal to 512 μg/ml. Each assay was performed at least 3 times. *** p<0.001; **** p<0.0001

The poor solubility of TS has hampered its development as a therapeutic, but these data suggested that its effective concentration could be reduced in the presence of iron chelators. We tested the FDA-approved iron chelators deferiprone (DFP) and deferasirox (DSX) for potential synergy with TS. Checkerboard assays revealed that while neither chelator had activity against *P. aeruginosa* on its own, both potentiated TS activity (**Fig 3DE**) at concentrations well below those used to safely treat patients, up to 28 mg/kg/day for DSX or 99 mg/kg/day for DFP (36).

### TS hijacks pyoverdine receptors FpvA and FpvB

To identify the route of iron-limitation dependent TS entry into *P. aeruginosa*, we tested the susceptibility of mutants from the ordered PA14 transposon library (37) that had insertions in genes encoding known siderophore receptors, as well as mutants with insertions in uncharacterized OM proteins with homology to siderophore receptors. In VBMM, most mutants had an IC_50_ of 0.16 μM (0.26 μg/ml), similar to the parental strain (**Table 1**). In contrast, an *fpvA* mutant, encoding the type I pyoverdine receptor, had an IC_50_ of 3.5 μM. Growth inhibition was still observed at the highest TS concentrations, indicating that the *fpvA* mutant remained partially susceptible. *P. aeruginosa* encodes two type I pyoverdine receptors, FpvA and FpvB, with ~39% amino acid identity (71% similarity). The *fpvB* mutant was also less susceptible to TS than the parent strain, with an IC_50_ of 1 μM. Based on these patterns of susceptibility, we speculated that TS may use both FpvA and FpvB, but that FpvA was the preferred receptor. When we deleted *fpvA* in the *fpvB* background, the double mutant was more resistant to TS than the single mutants, with an IC_50_ of 6.3 μM (**Table 1**). Complementation of the *fpvAB* mutant with *fpvB* on a plasmid restored TS susceptibility to levels similar to those of the single *fpvA* mutant (Table 1). Together, these data suggest that TS exploits both pyoverdine receptors for entry.

**Table 1.** Susceptibility of *P. aeruginosa* mutants to thiostrepton in VBMM.

| Strain | Gene | Function | IC<sub>50</sub> µg/ml | IC<sub>50</sub> µM |
| --- | --- | --- | --- | --- |
| PA14 | N/A | Wild type strain | 0.26 | 0.16 |
| *fpvA*::Mar2xT7 | PA2398 | Type 1 ferripyoverdine receptor | 3.68 | 2.21 |
| *fpvB*::Mar2xT7 | PA4168 | Alternate Type 1 ferripyoverdine receptor | 1.83 | 1.10 |
| *pvdA*::Mar2xT7 | PA2386 | L-ornithine N5-oxygenase | 0.26 | 0.16 |
| Δ*fpvA*<br>*fpvB*::Mar2xT7 | PA2398,<br>PA4168 | Type I ferripyoverdine receptor,<br>Alternate type I ferripyoverdine receptor | 6.93 | 4.16 |
| Δ*fpvA*<br>*fpvB*::Mar2xT7 +<br>pUCP20 | PA2398,<br>PA4168 | Type I ferripyoverdine receptor,<br>Alternate type I ferripyoverdine receptor | 4.20 | 2.52 |
| Δ*fpvA*<br>*fpvB*::Mar2xT7 +<br>pUCP20-*fpvB* | PA2398,<br>PA4168 | Type I ferripyoverdine receptor,<br>Alternate type I ferripyoverdine receptor | 2.80 | 1.68 |
| *fptA*::Mar2xT7 | PA4221 | Pyochelin receptor | 0.26 | 0.16 |
| PA1322::Mar2xT7 | PA1322 | Probable TonB-dependent receptor | 0.26 | 0.16 |
| PA4837::Mar2xT7 | PA4837 | Probable outer membrane protein precursor | 0.26 | 0.16 |
| *foxA*::Mar2xT7 | PA2466 | Ferrioxamine receptor FoxA | 0.26 | 0.16 |
| *fiuA*::Mar2xT7 | PA0470 | Ferrichrome receptor FiuA | 0.26 | 0.16 |
| *pirA*::Mar2xT7 | PA0931 | Alternate enterobactin receptor | 0.35 | 0.21 |
| *pfeA*::Mar2xT7 | PA2688 | Enterobactin receptor | 0.26 | 0.16 |
| *hasR*::Mar2xT7 | PA3408 | Heme uptake outer membrane receptor | 0.35 | 0.21 |
| *fvbA*::Mar2xT7 | PA4516 | Vibriobactin receptor | 0.26 | 0.16 |
| *piuA*::Mar2xT7 | PA4514 | Hydroxamate-type ferrisiderophore receptor | 0.35 | 0.21 |
| PA0151::Mar2xT7 | PA0151 | Probable TonB-dependent receptor | 0.26 | 0.16 |
| chtA::Mar2xT7 | PA4675 | Aerobactin, Rhizobactin 1021,<br>Schizokinen receptor | 0.59 | 0.36 |
| phuR::Mar2xT7 | PA4710 | Heme/Hemoglobin uptake receptor<br>precursor | 0.26 | 0.16 |

### TS is active against multi-drug resistant clinical isolates

To test whether TS could inhibit growth of a broader range of *P. aeruginosa* strains, particularly multi-drug resistant (MDR) isolates for which there are fewer antibiotic options, we tested 96 recent clinical isolates for susceptibility to TS in 10:90. While approximately 1 in 10 of those strains had an IC_50_ ≥5μM TS (**Fig 4A**), a combination of 5 μM TS (8.3 μg/ml) plus 86 μM DSX (32 μg/ml) reduced growth of all but three isolates of *P. aeruginosa* to less than 25% of the DMSO control (**Fig 4A**). We next tested the activity of TS against another MDR Gram-negative pathogen that can cause severe infections, *Acinetobacter baumanii* (38). *A. baumannii* strains encode FpvA and FpvB homologs (**Supplementary Fig S3**), suggesting they may be susceptible to the thiopeptide. Growth of 6 of 10 *A. baumannii* strains in 10:90 was reduced to ≤50% of control with 5 μM TS, while the combination of 5μM TS and 86 μM DSX reduced growth of 8/9 clinical isolates of *A. baumanii* below 10% of control (**Fig 4B**). As reported previously (39), growth of *E. coli*– which lacks FpvAB homologs – was unaffected even at the maximum soluble concentration of 10 μM TS (not shown).

**Figure 4.**
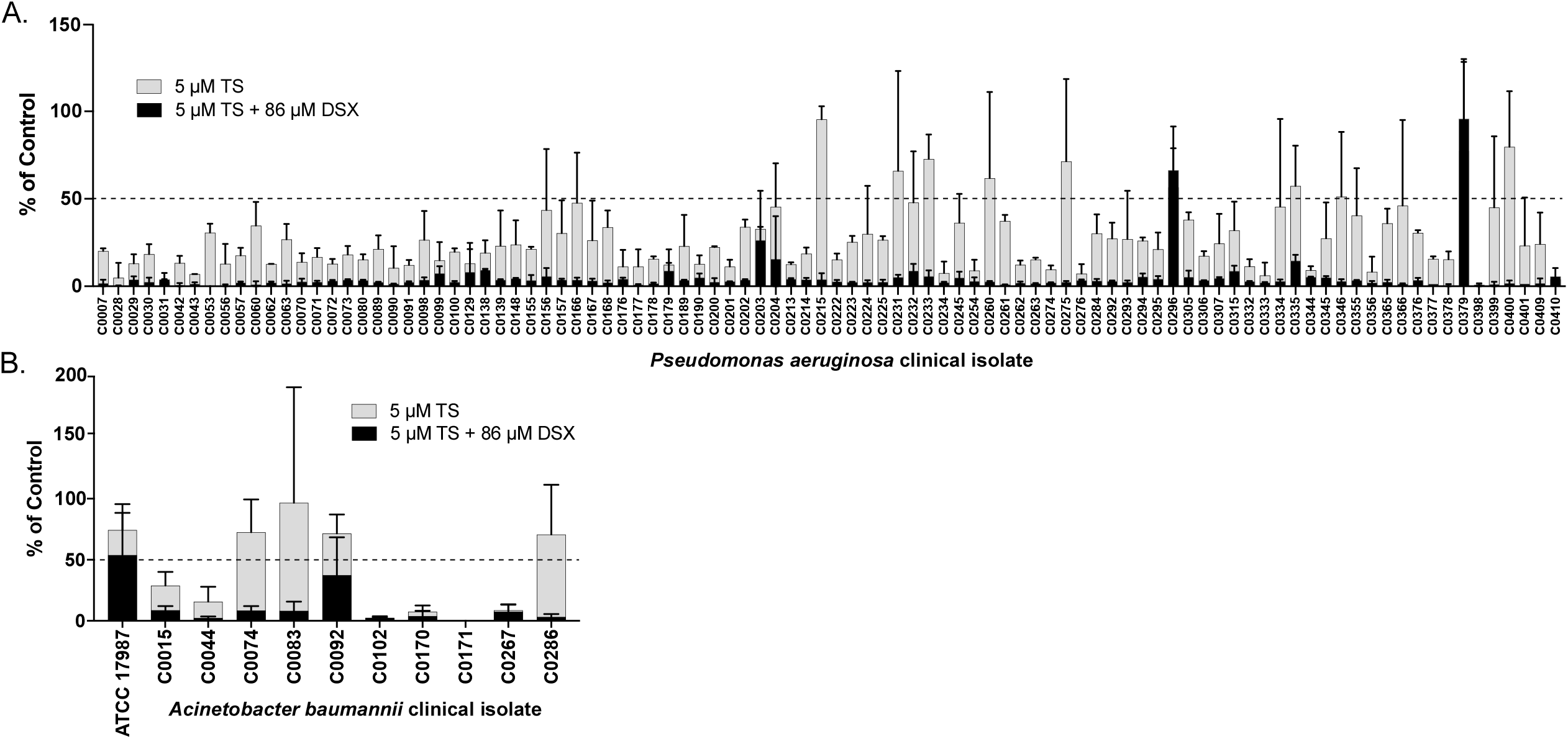
Thiostrepton inhibits growth of multidrug-resistant clinical isolates. The growth of most clinical isolates of A. *P. aeruginosa* and B. *Acinetobacter baumannii* resistant to multiple antibiotics (see **Supplementary Table S2** for antibiograms) was inhibited by 5 μM (8.3 μg/ml) TS in 10:90 medium (grey bars). TS activity was potentiated by the addition of 86 μM deferasirox (DSX; 32 μg/ml; black bars). Each assay was performed at least 3 times and the results plotted as percent of the DMSO-only growth control (OD_600_) using Prism (GraphPad). Error bars equal one standard deviation.

## DISCUSSION

The natural role of antibiotics has been broadly debated (14, 16): are they signaling molecules that are toxic at high concentrations, or weapons used by bacteria to gain an advantage over competitors in their environment? The biofilm stimulation response to sub-inhibitory concentrations of antibiotics is consistent with both views. At concentrations too low to elicit damage, bacteria show little phenotypic response to antibiotic exposure. As concentrations approach the MIC, the bacteria respond in a dose-dependent manner by ramping up the amount of biofilm produced – detecting either the antibiotics or their effects – which may protect a subpopulation of cells. Above the MIC, antibiotics fall into the deadly weapons category. Biofilm stimulation by sub-inhibitory concentrations of antibiotics is a common phenomenon among multiple gram-positive and gram-negative species, and is caused by several drug classes, suggesting it is not necessarily linked to a specific MOA (16, 18, 40). As demonstrated here, this phenomenon can be used to identify potential antibiotic activity in the absence of killing, a useful feature when screening at a single arbitrary concentration that may be below the MIC for the drug-organism combination being tested. Interestingly, we and others (41) found that many drugs intended for eukaryotic targets can impact bacterial growth and biofilm formation (**Supplementary Table S1**), implying that they have deleterious effects on prokaryotic physiology. With a new appreciation of the role of the human microbiome in health and disease, these potential effects should be considered during drug development.

TS, a complex cyclic thiopeptide made by *Streptomyces azureus, S. hawaiiensis*, and *S. laurantii*, is experiencing a resurgence of research interest due to its broad anti-bacterial, anti-malarial, and anti-cancer activities (23, 24). It is a member of the RiPP (ribosomally synthesized and post-translationally modified peptides) class of natural products (42), derived from a 42-amino acid precursor, TsrA (43). Although the mechanism of its antibacterial activity (inhibition of translation by binding to helices H43/H44 of 23S rRNA) and resistance (methylation of 23S rRNA residue A1067) have been deciphered (26, 44), the way in which this ~1.7 kDa molecule enters target bacteria is unknown. Our data suggest that TS is actively imported into *P. aeruginosa* under iron-restricted conditions.

There are multiple examples of molecules that exploit iron uptake pathways to enter bacteria. Class I microcins – narrow-spectrum antibiotics produced by some gram negative species – bind to siderophore receptors and share many of TS’s properties. They are RiPPs, less than 5 kDa in mass, and cyclic (giving them the nickname ‘lasso peptides’). Notably, binding of iron by microcins is not a prerequisite for uptake, as some interact with siderophore receptors in an iron-free state. For example, MccJ25, produced by *E. coli* (45), interacts with siderophore receptor FhuA by mimicking the structure of ferrichrome (46). It is not yet clear whether TS binds iron or simply mimics an iron-bound conformation, as it has multiple hydroxyls positioned in a manner that could coordinate metals (**Fig 1A**). Its structure has been solved both by X-ray crystallography and NMR, but no bound metals were reported (47, 48). The FpvA receptor is also exploited by S-pyocins, 40-80 kDa peptide antibiotics produced by competing *P. aeruginosa* strains, showing that it is a key promiscuous entry point for diverse molecules in addition to its usual ligand, pyoverdine (33, 34, 49).

Our discovery that TS exploits FpvA and FpvB for uptake into the periplasm helps to explain the resistance of gram-negative species such as *E. coli* to this antibiotic, as they lack those proteins. Bioinformatic searches show FpvAB homologs are expressed by *P. aeruginosa* and related pathogens – including *A. baumanii* (**Supplementary Fig S3**) – suggesting that TS could have utility as a narrow-spectrum agent. Use of multiple pyoverdine receptors by TS may reduce the probability of resistance arising through mutation of a single receptor, although genome analysis of clinical isolate C0379 that was most resistant to the combination of TS and DSX (**Fig 4A**) revealed a wild type copy of *fpvA* coupled with an ~800 bp deletion encompassing the 5’ region of *fpvB*. Further investigation of *P. aeruginosa* strains that are resistant to TS will be needed to understand the most likely routes by which it occurs.

Although TS uses siderophore receptors to cross the *P. aeruginosa* OM, the way in which this large cyclic peptide transits the cytoplasmic membranes of gram-positive and gram-negative bacteria to reach its ribosomal targets remains undefined. Expression of *tsr* in *P. aeruginosa* conferred resistance, confirming that TS acts via its canonical bacteriostatic MOA. While PA14 expressing Tsr was significantly more resistant to TS than the control, it was more sensitive than PAO1. This difference is not due to nucleotide polymorphism at the Tsr methylation site on the rRNA, as these residues are conserved between PAO1 and PA14. The reasons for strain-specific differences in susceptibility are unclear, but our data confirm that most *P. aeruginosa* isolates tested (including MDR strains) are susceptible to TS, especially when it is combined with DSX (**Fig 4A, Supplementary Table S2**).

TS’s major liability is its poor solubility (50). Smaller, more soluble fragments that retain activity against gram-positive bacteria and have reduced toxicity for eukaryotic cells have been identified (51) but it is not clear if they would be active against *P. aeruginosa* if uptake by the FpvAB receptors requires the intact molecule. Another way to circumvent solubility issues is to reduce the required concentration. Our data show that this can be accomplished for TS by co-administration with FDA-approved iron chelators DFP or DSX (**Fig 3DE**). The true potential of TS as an anti-infective may be underestimated, as MIC evaluations are typically performed in rich, iron-replete media such as cation-adjusted Mueller-Hinton broth. Many host environments are iron-restricted, particularly in the presence of infection and inflammation (52–54) and future studies of TS activity should emulate those growth conditions.

In summary, we showed that biofilm stimulation can be used in high-throughput small molecule screening to report on sub-inhibitory antibiotic activity that may otherwise be missed using the usual metric of growth inhibition. In a small screen of less than 4000 molecules at a fixed concentration of 10 μM, we identified 60 molecules that stimulated biofilm formation, suggesting that they may have antimicrobial activity at higher concentrations, or under different growth conditions, as demonstrated here for TS. Stimulation of biofilm matrix production by TS in the gram-positive genus *Bacillus* was reported previously, and that phenotype leveraged to identify novel thiopeptide producers in co-cultures (55). Those studies, and the data presented here, suggest that monitoring biofilm stimulation (or an easily assayed proxy thereof, such as increased expression from biofilm matrix promoters) could allow for increased detection of molecules with potential antibacterial activity during screening, making it a useful addition to the antimicrobial discovery toolkit.

## ACKNOWLEDGEMENTS

We thank Gerry Wright for access to strains from the Wright Clinical Collection, David Heinrichs for the gift of EDDHA and helpful discussions, and Neha Sharma, Andrew Hogan, Amanda Veri, and Victor Yang for assistance with method development. This work was supported by a Natural Sciences and Engineering Research (NSERC) grant RGPIN-2016-06521, and by Ontario Research Fund grant RE07-048. MRR and UN held Ontario Graduate Scholarships, MR was supported by an NSERC Undergraduate Summer Research Award, SKP was supported by a Summer Studentship from GlycoNet, and HA was supported by a Summer Studentship from Cystic Fibrosis Canada.

## METHODS

### Bacterial strains and culture conditions

The bacterial strains and plasmids used in this study are listed in **Table 1** and **Supplementary Table S2**. Bacterial cultures were grown in Lysogeny Broth (LB), 10:90 (10% LB and 90% phosphate buffered saline), M9 medium, Vogel Bonner minimal medium (VBMM), or cation-adjusted Mueller-Hinton broth (MBH) as indicated. Where solid media were used, plates were solidified with 1.5% agar. DFP (Sigma-Aldrich) and DSX (Cayman Chemicals) were stored at 4°C until use. TS was stored at °20°C. A 60 mg/mL stock solution of DFP was made in 6M HCl and Milli-Q H2O (DFP solvent) in a ratio of 3:50. A 20 mg/mL stock solution of DSX was made in DMSO. A 20mM stock solution of TS was made in DMSO.

### Growth curves

PAO1-KP was inoculated from a −80°C stock into 5 ml LB broth and grown with shaking at 200 rpm, 16h, 37°C. The overnight culture was subcultured at 1:500 into 5 different media (LB, 10:90, M9, Mueller-Hinton (MH), and VBMM) – incubated at 37°C for 6h with shaking at 200 rpm. Each subculture was standardized to OD_600_ ~ 0.1 (Biomate 3 Spectrophotometer) then diluted 1:500 into the same medium. Six replicates of 200 μl of each sample were added to a 96 well plate, which was incubated at 37°C for 24 h with shaking at 200 rpm (Tecan Ultra Evolution plate reader). The OD612 was read every 15 min for 24 h. The data for the six replicates of each sample were averaged and the experiment was repeated 3 times. The final data with standard deviations were plotted using Prism (Graphpad).

### Biofilm modulation assay

Biofilm formation was assayed as described in (11), with modifications. Briefly, *P. aeruginosa* was inoculated in 5 mL of LB and grown at 37°C overnight, shaking at 200 rpm, and subsequently standardized to an OD_600_ of ~ 0.1 in 10:90. For the initial screen, 1 mM compound stocks in DMSO were diluted 1:100 in standardized cell suspension (1.5 μL of compound stock in 148.5 μL of cell suspension) to a final concentration of 10 μM. Control wells contained 10:90 plus 1% DMSO (sterility control) or standardized cell suspension plus 1% DMSO (growth control). Biofilms were formed on polystyrene peg lids (Nunc). After placement of the peg lid, the plate was sealed with parafilm to prevent evaporation and incubated for 16 h at 37°C, 200 rpm. Following incubation, the 96-peg lid was removed and planktonic density in the 96 well plate measured at OD_600_ to assess the effect of test compounds on bacterial growth. The lid was transferred to a new microtiter plate containing 200 μl of 1X phosphate-buffered saline (PBS) per well for 10 min to wash off any loosely adherent bacterial cells, then to a microtiter plate containing 200 μL of 0.1% (wt/vol) CV per well for 15 min. Following staining, the lid was washed with 70 mL of dH2O, in a single well tray, for 10 min. This step was repeated four times to ensure complete removal of excess CV. The lid was transferred to a 96-well plate containing 200 μL of 33% (vol/vol) acetic acid per well for 5 min to elute the bound CV. The absorbance of the eluted CV was measured at 600 nm (BioTek ELx800), and the results plotted as percent of the DMSO control using Prism (Graphpad). Screens were performed in duplicate. Compounds that resulted in <50% of control biofilm were defined as biofilm inhibitors, while compounds that resulted in >200% of control biofilm were defined as biofilm stimulators. Compounds of interest were further evaluated using the same assay but over a wider range of concentrations (dose-response assay).

For TS dose response assays, TS stock solutions were diluted in DMSO and 2μL of the resulting solutions plus 148μL of a bacterial suspension standardized to an OD_600_ of ~ 0.1 in 10:90 were added to a 96 well plate in triplicate, as described above. Control wells contained 148μL of 10:90 + 1.3% DMSO (sterility control) or standardized bacterial suspension + 1.3% DMSO (growth control). For EDDHA alone or with FeCl_3_ experiments, 2μL of each were added as aqueous solutions to reach final concentrations of 0.1μM EDDHA and 100μM FeCl_3_, and the amount of bacterial suspension adjusted to keep the total well volume at 150μL. Controls for EDDHA and FeCl_3_ were 2μL of sterile dH20. Biofilms were grown for 16h at 37°C, 200 rpm, then stained and quantified as described above. Assays were performed in triplicate and results were graphed using Prism (Graphpad) as a percentage of the DMSO control.

### Compounds screened

The biofilm modulation assay was used to screen the McMaster Bioactives compound collection. This curated collection includes off-patent, FDA-approved drugs from the Prestwick Chemical Library (Prestwick Chemical, Illkirch, France), purified natural products from the Screen-Well Natural Products Library (Enzo Life Sciences, Inc., Farmingdale, NY, USA), drug-like molecules from the Lopac^1280^ (International Version) collection (Sigma-Aldrich Canada Ltd., Oakville, ON, Canada) and the Spectrum Collection (MicroSource Discovery Systems, Inc., Gaylordsville, CT, USA) which includes off patent drugs, natural products, and other biologically active compounds. In total, the collection is 3921 unique compounds.

### Construction of a *tsr* plasmid for expression in *P. aeruginosa*

The *tsr* gene from pIJ6902 (56) was PCR-amplified using primers 5’ GAATCCCGGGCGGTAGGACGACCATGAC 3’ and 5’ CTTCAAGCTTTTATCGGTTGGCCGCGAG 3’. Both the PCR product and pUCP20 vector were digested with SmaI and HindIII, gel-purified, and ligated at a 1:3 molar ratio using T4 DNA ligase. The ligated DNA was transformed into *E. coli* DH5a and transformants selected on LB agar containing 100 μg/mL ampicillin and 5-bromo-4-chloro-3-indolyl-β-D-galactopyranoside for blue-white selection. Plasmids from white colonies were purified using a GeneJet Plasmid Miniprep kit (Thermo Scientific) following the manufacturer’s protocols. After verification by restriction digest and DNA sequencing, pUCP20 and pUCP20-tsr were each introduced into *P. aeruginosa* PAO1 and PA14 by electroporation. Transformants were selected on LB agar containing 200 μg/mL carbenicillin.

### IC_50_ and checkboard assays

IC_50_ values were determined with microbroth dilution assays in Nunc 96-well plates. Vehicle controls consisted of 1:75 dilutions of DMSO in 10:90 inoculated with PA14 or its mutants as described in Growth Curves. Sterile controls consisted of 1:75 dilutions of DMSO in 10:90. Seven serially diluted concentrations of TS – with 17 μg/mL being the highest final concentration – was set up in triplicate. Tests were done with 1:75 dilutions of each TS concentration in 10:90 inoculated with PA14 or its mutants as described in Growth Curves. Plates were sealed to prevent evaporation and incubated with shaking at 200 rpm, 16h, 37°C. The OD_600_ of the plates was read (Multiskan Go - Thermo Fisher Scientific) and used to calculate IC_50_. The final volume of each well was 150 μL and each experiment was repeated at least three times.

Checkerboard assays were set-up using Nunc 96-well plates in an 8-well by 8-well format. Two columns were allocated for vehicle controls and two columns for sterility controls. Vehicle controls contained 2 μL DMSO + 2 μL DFP solvent for checkerboards with TS and DFP or 4 μL DMSO for TS and DSX, in 146 μL of 10:90 inoculated with PA14 or PAO1-KP as described in Growth Curves. Sterile controls contained the same components in 10:90, without cells. Serial dilutions of TS – with 17 μg/mL being the highest final concentration – was added along the ordinate of the checkerboard (increasing concentration from bottom to top) whereas serial dilutions of DFP or DSX – with 512 μg/mL being the highest final concentration – was added along the abscissa (increasing concentration from left to right). The final volume of each well was 150 μL and each checkerboard was repeated at least three times. Plates were incubated and the final OD_600_ determined as detailed above.

### Clinical isolates testing

Clinical isolates of *P. aeruginosa* and *A. baumannii* were inoculated from −80°C stocks into 200 μL LB broth and grown with shaking at 200 rpm, 16h, 37°C. in Nunc 96-well plates. The overnight cultures were subcultured (1:25 dilution) into 10:90 and grown with shaking at 200 rpm, 2h, 37°C. Vehicle controls consisted of 4 μL of DMSO, 144μL 10:90 and 2 μL of subculture. Sterile controls consisted of 4 μL of DMSO and 146μL 10:90. Test samples consisted of 2 μL of TS (final concentration of 5 μM, 8.3 μg/ml), 2 μL of DMSO (or DSX, final concentration of 86 μM, 32 μg/mL), 144 μL 10:90 and 2 μL of subculture. The final volume of each well was 150 μL and each checkerboard was repeated at least three times. Plates were incubated with shaking at 200 rpm, 16h, 37°C and OD_600_ was measured (Multiskan Go - Thermo Fisher Scientific). The results were plotted as percent of control (wells containing only DMSO) using Prism (GraphPad).

